# *SIM* and *CCS52A1* control root hair cell endoreplication and expansion

**DOI:** 10.1101/2024.10.20.618934

**Authors:** Lei Li (李磊), Klaus Herburger, Ilse Vercauteren, Jefri Heyman, Lieven De Veylder

## Abstract

*Arabidopsis thaliana* (Arabidopsis) root hair cells undergo rapid cell elongation that is correlated with an increased DNA ploidy level through endoreplication. However, the contribution of endoreplication to hair cell growth is currently unclear. Here, we determined ploidy levels within the root hair cell file from the proximal meristem to the elongation zone and correlated them with cellular parameters. Our data show that cells in the basal meristem reach a 4C level prior to rapid elongation, whereas in the elongation zone a 16C level is reached prior to the onset of tip growth, suggesting that endoreplication is a prerequisite for rapid volume increase. Consistently, we show that mutants in *SIAMESE* (*SIM*) and *CELL CYCLE SWITCH52 A1* (*CCS52A1*), key endocycle regulators expressed in hair cells, display a decrease in cell volume and hair tip length. Interestingly, SIM and CCS52A1 affect cell width and length, respectively. In addition, CCS52A1 controls the growth rate of the root hair tip. Given the common association between cell expansion and cell wall modifications, we performed immunolabeling on wild-type and mutant roots, which revealed a reduced level of pectin methylesterification in the mutant backgrounds, suggesting that SIM and CCS52A1 may modulate root hair cell growth by affecting cell wall modifications.

The author responsible for distribution of materials integral to the findings presented in this article in accordance with the policy described in the Instructions for Authors (https://academic.oup.com/plphys/pages/General-Instructions) is Lieven De Veylder.

**ONE-SENTENCE SUMMARY:** *SIM* and *CCS52A1* control Arabidopsis root hair cell volume by controlling endoreplication.

## Introduction

The root is a vital organ supporting plant growth and development by soil anchorage and uptake of water and nutrients (Dietrich, 2018; Dinneny, 2019). Root growth displays significant plasticity, determined by the availability and distribution of water and nutrients in the soil (Roy and Bassham, 2014; Karlova et al., 2021; Zou et al., 2022). Primary root growth is driven by the root tip that consists of three main regions: the meristematic zone, the elongation zone (EZ), and the differentiation zone (Petricka et al., 2012; Motte et al., 2019). Within the meristematic zone, the root cells divide to ultimately expand in the EZ, where anisotropic expansion dominates. The boundary between both is known as the transition zone (TZ), where root epidermal cells undergo rapid elongation (Takatsuka and Umeda, 2015).

The root stem cells surrounding the quiescent center (QC) cells in the root meristematic zone give rise to the diverse tissue types, including the epidermis, cortex, endodermis, columella, lateral root cap, pericycle, phloem, procambium, metaxylem, and protoxylem (Matosevich and Efroni, 2021; Strotmann and Stahl, 2021). Epidermal cells, originating from epidermis/lateral root cap stem cells, can be classified into trichoblast [hair (H)] and atrichoblast [non-hair (N) cells]. H cells are located at the junction between two underlying cortical (C) cells, while N cells arise from a single C cell (Grierson et al., 2014; Bienert et al., 2021). Typically, the young Arabidopsis primary root holds 8 H and 10-14 N cell files. Multiple factors driving root H and N cell specification have been identified (Grierson et al., 2014; Shibata and Sugimoto, 2019), with *CAPRICE* (*CPC*), *TRIPTYCHON* (*TRY*), and *ENHANCER OF TRY AND CPC* (*ETC1*) promoting H cell file specification, whereas *TRANSPARENT TESTA GLABRA* (*TTG*), *GLABRA3* (*GL3*), *ENHANCER OF GLABRA3* (*EGL3*), and *WEREWOLF* (*WER*) establish the formation of the N cell files (Chen and Schmidt, 2015; Li et al., 2015; Tominaga-Wada and Wada, 2016; Wang et al., 2019).

In the meristematic zone, there are no apparent morphological differences between H and N cell files. In the root differentiation zone, root hairs develop on the soil-facing surface of H cells as tubular-like structures, driven by tip growth. Tip growth of H cells entails a specific type of polarized cell expansion that is precisely regulated by various factors, including Ca^2+^ signaling, reactive oxygen species, cell wall modifications, microtubule and actin organization, and cellular pH (Wang et al., 2015; Dindas et al., 2018; Gayomba and Muday, 2020; Bienert et al., 2021; Feiguelman et al., 2022). Additionally, tip growth is intricately controlled by multiple plant hormones, each employing distinct mechanisms (Li et al., 2022). For instance, auxin plays a pivotal role by triggering calcium signaling during root hair tip extension (Dindas et al., 2018), while cytokinin has been found to promote root hair elongation via activating the *ROOT HAIR DEFECTIVE 6-LIKE4* (*RSL4*) transcription factor (Takatsuka et al., 2023). Environmental factors, such as water content, soil pH value and heavy metal ions, also profoundly influence root hair growth (Bienert et al., 2021; Stéger and Palmgren, 2022).

Previous work has identified the root TZ as the site of the mitotic-to-endoreplication cycle switch (Bhosale et al., 2018). Endoreplication is a specialized cell cycle, widespread in higher plants, mammals, and arthropods (Qi and Zhang, 2020; Shimotohno et al., 2021), in which the nuclear genome replicates but mitosis is skipped (Breuer et al., 2010; Edgar et al., 2014). In plants, endoreplication has been implicated in establishing trichome size, the hypocotyl length, and root meristem size (Vanstraelen et al., 2009; Kumar et al., 2015; Ma et al., 2022; Goldy et al., 2023). The onset of endoreplication has been demonstrated to be associated with a shift from high to low cyclin-dependent kinase (CDK) activity (De Veylder et al., 2011), which can be achieved through either the inhibition of CDK activity or the degradation of their regulatory cyclins partners. The SIAMESE/SIAMESE-RELATED (SIM/SMR) family represents a group of CDK inhibitors that has been proven to promote the onset of endoreplication (Walker et al., 2000; Churchman et al., 2006). The Arabidopsis *sim* loss-of-function mutant develops multicellular trichomes with a lower ploidy level than the normal single-cell structures observed in wild-type (WT) plants (Churchman et al., 2006). Moreover, *SIM*, *SMR1* (also known as *LOSS OF GIANT CELLS*), and *SMR2* function redundantly in controlling endocycle onset in Arabidopsis leaves (Kumar et al., 2015). Overexpression of *SMR1* results in giant cells with higher ploidy levels in *Arabidopsis* sepals (Schwarz and Roeder, 2016). Another pathway triggering endoreplication involves the degradation of cyclins by the anaphase-promoting complex/cyclosome (APC/C), a conserved E3 ubiquitin ligase. The Arabidopsis CELL CYCLE SWITCH 52 A1 (CCS52A1) and A2 (CCS52A2) proteins are activating subunits of the APC/C and control endocycle onset by promoting cyclin protein ubiquitination that triggers their proteolytic turn-over (Willems and De Veylder, 2022). As such, *ccs52a1* and *ccs52a2* knock-outs exhibit endoreplication onset defects (Vanstraelen et al., 2009; Breuer et al., 2010; Breuer et al., 2012; Liu et al., 2012). Corresponding with a role for both *SIM* and *CCS52A1* in controlling endocycle onset in the Arabidopsis root, both genes are reported to be expressed in the root TZ that represents the boundary between the mitotic cycle and the endocycle (Ishida et al., 2010; Takahashi et al., 2013; Bhosale et al., 2018).

Endoreplication is typically associated with cell expansion, in plants illustrated by a frequently observed correlation between epidermal cell size and the ploidy level (Bhosale et al., 2019). Accordingly, through mathematical modeling, it was reported that Arabidopsis H cells undergo an additional round of endoreplication compared to N cells (Bhosale et al., 2018), suggesting a role for the endocycle in promoting root tip growth. However, the correlation between ploidy level and cell size may not be causal, because many cell types can still enlarge in the absence of endoreplication (Sliwinska et al., 2015; Tsukaya, 2019). Accordingly, whereas *midget* and *rhl1/hyp7* mutants, which have an endocycle defect due to a mutation in the DNA topoisomerase VI complex, display impaired H cell growth (Sugimoto-Shirasu et al., 2005; Kirik et al., 2007), there appears to be no correlation between collet hair cell length and their ploidy level.

In recent years, the association of cell wall composition with endoreplication status has attracted increasing attention (Bhosale et al., 2019). Previous studies have found that homogalacturonan (HG), a major component of the primary cell wall, is synthesized in a highly methylesterified form before it is incorporated into the growing cell wall, followed by enzymatic demethylation via pectin methyl-esterases (PMEs). Thus, this process results in an increase of demethylesterified HG during cell expansion (Kohorn, 2016; Saffer, 2018; Haas et al., 2020; Shin et al., 2021). Interestingly, immunolabeling of root hair tips using cell wall-directed antibodies revealed significant cell wall modifications during root hair elongation (Herburger et al., 2022), whereas a recent study revealed that the mutation in *CCS52A2*, a homolog of *CCS52A1*, enhanced the methylesterification level of HG and inhibited endoreplication in the apical hook of dark-grown Arabidopsis seedlings, resulting in impaired cell expansion (Ma et al., 2022). These findings imply that endoreplication may regulate cell expansion via cell wall modifications.

In this study, we aimed to elucidate the function of SIM and CCS52A1 in root hair cell expansion. Through DNA ploidy mapping and cellular annotation using MorphoGraphX (MGX) software (Strauss et al., 2022), we assessed the endoploidy levels in the roots of WT, *sim* and *ccs52a1* single and *sim ccs52a1* double mutants. Immunolabeling of the cell wall indicated significant changes in cell wall modifications in root hairs of *sim*, *ccs52a1*, and *sim ccs52a1* mutants compared to the control. These findings potentially explain the observed short length of mutant root hairs and indicate that *SIM* and *CCS52A1* may regulate H cell expansion by modulating cell wall modifications during the endoreplication cycle.

## Results

### Hair cell expansion is preceded by a ploidy increase

Firstly, we aimed to investigate how epidermal cell volumes changes along the root tip. Utilizing the MGX ‘3DCellAtlas’ function, a systematic grouping of root epidermal cells was performed based on their relative position along the root axis (Strauss et al., 2022). The position of the QC was set as the 0.0 point, and the fifth/sixth elongated cell following onset of the TZ as the 1.0 point (Fig. 1A). Cell volumes in both H and N cell files displayed a notable increase in cell volume in the region between 0.4 and 0.6 (Fig. 1B), marking it as the TZ. Accordingly, we categorized the epidermal cells into four subgroups according to their location: the proximal meristem (PM, 0.0 to 0.2), the basal meristem (BM, 0.2 to 0.4), the TZ (0.4 to 0.6), and the EZ (0.6 to 1.0). To discern the differences between H and N cells, we compared their cell volume, length, and width. Results revealed a significantly larger volume of H cells compared to N cells in the meristematic zones (PM and BM), but no significant differences were seen within the TZ and EZ (Figs. 1C-F). Geometrical analysis illustrated that cell length and width of H cells exceeded those of N cells in the meristematic zones (Supplementary Fig. S1), contributing to the observed higher cellular volume. Within the TZ and EZ, N cells became longer than H cells, while H cells maintained a higher width than N cells (Supplementary Fig. S1), resulting in two distinct morphological identities.

**Fig. 1.**
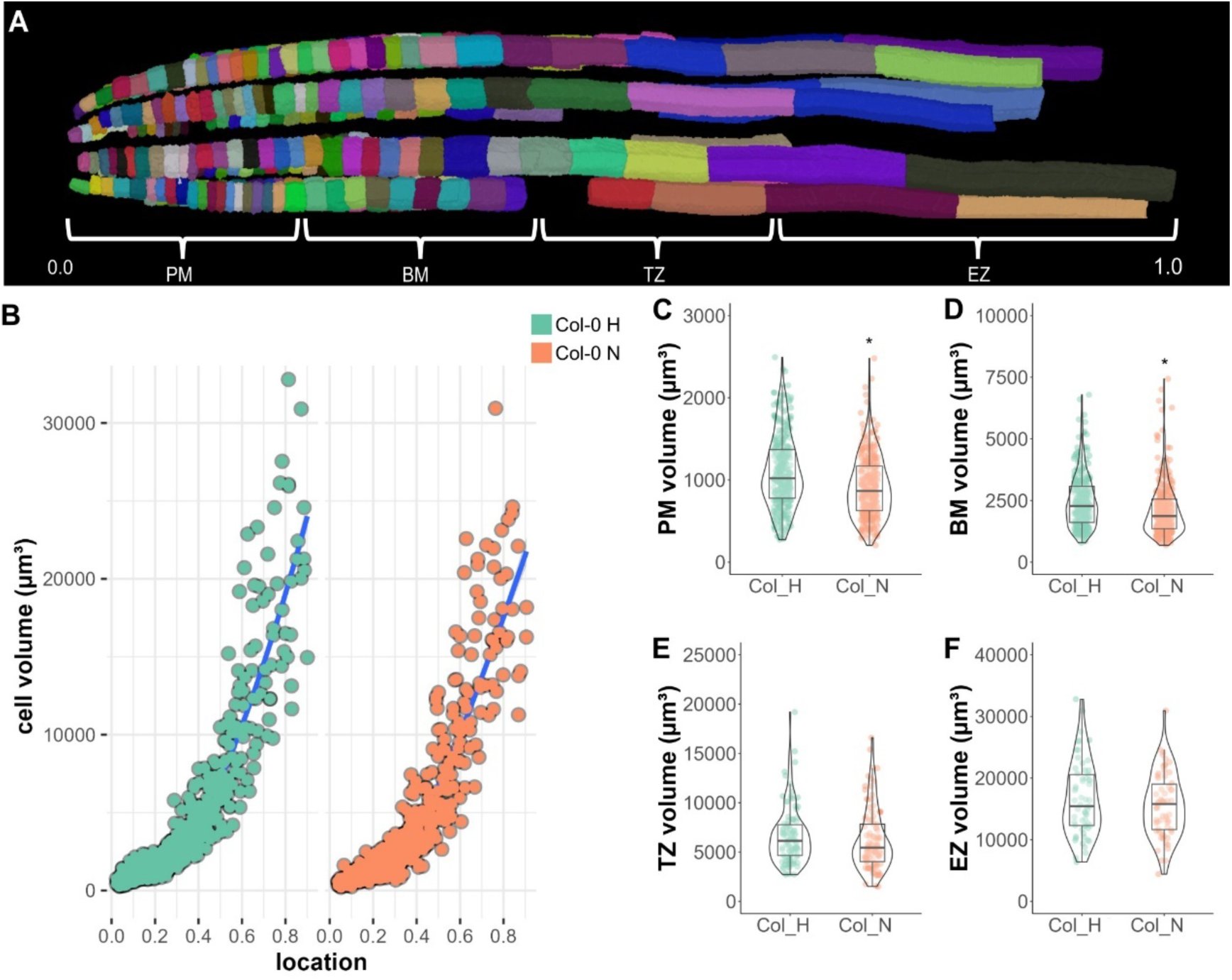
Comparison of cell volume between H and N cells in the Arabidopsis primary root. **A**, H cells in the primary root from the QC to the fifth/sixth elongated cell defined as 0.0 and 1.0 points, respectively. According to their location along the root longitudinal direction, the epidermal cells were categorized into four zones, being the proximal meristem (PM), basal meristem (BM), transition zone (TZ), and elongation zone (EZ). **B**, The H and N cell volume distribution along the root longitudinal axis. Blue lines show Loess regression curves. Over 600 cells from 3 independent roots for each cell type were analyzed. **C-F**, Cell volumes of H and N cells within the PM (C), BM (D), TZ (E), and EZ (F). Asterisks indicate a significant difference (adjusted p < 0.05) determined by parametric t-test and post hoc Tukey test (n ≥ 60 cells per genotype).

Our prior mathematical modeling of DNA ploidy levels along the root axis predicted that H and N cells achieve ploidy levels of 16C and 8C, respectively (Bhosale et al., 2018). Using DAPI staining, we quantified the cellular DNA ploidy level in the WT from the 13 last cells within the BM up to the 8 to 9 earliest cells after leaving the meristematic zone, covering the complete TZ. In both the BM H and N cells, the predominant ploidy portion was 4C, with only the two first cells in the N cell files displaying a 2C DNA content, suggesting that cells in the BM already entered the endocycle (Fig. 2A and Supplementary Fig. S2). Within the TZ, cells in both cell files underwent an extra endocycle, reaching 8C. Within the EZ, H cells reached a 16C ploidy content, whereas N cells remained at 8C, confirming previous predictions (Bhosale et al., 2018). To validate this finding, we constructed a marker line with an H cell-specific promoter driving a histone *H2A-GFP* reporter gene (*AT2G34910:H2A-GFP*, Supplementary Fig. S3). H2A-GFP fluorescence preceded the strong increase in cell volume (Fig. 2B), but predominantly marked cells of the TZ and EZ. Accordingly, dual-color flow cytometry on the primary root confirmed the DNA ploidy levels of the GFP-positive cells within the elongating H cells to be 8C or 16C (Fig. 2C).

**Fig. 2.**
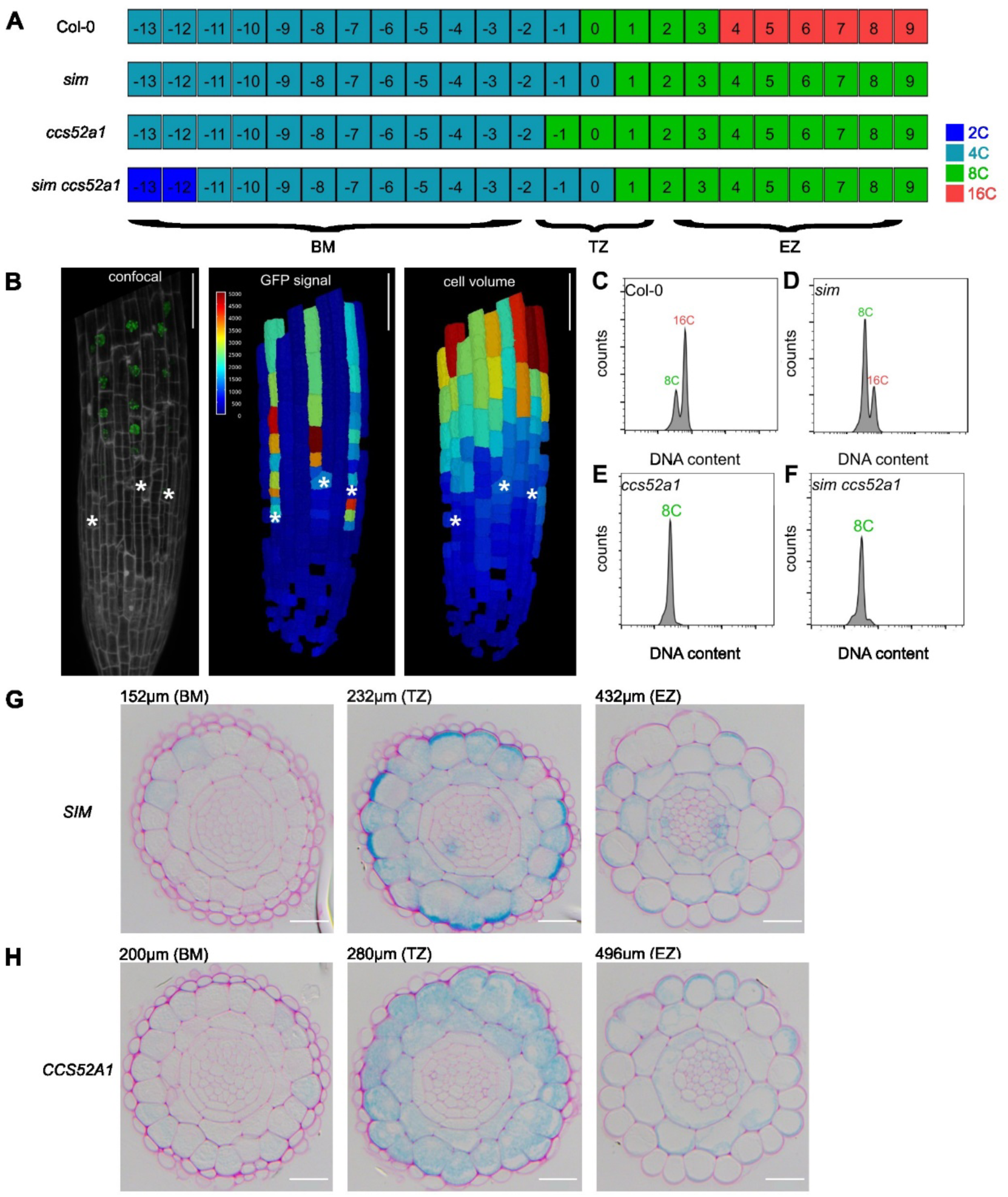
*SIM* and *CCS52A1* determine the endoreplication level of H cells in the Arabidopsis root. **A**, Mapping of H cell DNA ploidy levels along the root longitudinal axis of WT (Col-0) and *sim*, *ccs52a1* and *sim ccs52a1* mutants based on DAPI staining. The first elongated cell is set as the “0” point. Data were obtained from 4 to 5 independent roots. **B**, Representative confocal image of a *AT2G34910:H2A-GFP* root counterstained with propidium iodide (left), GFP expression heat map (middle), and cell volume heat map (right). MorphoGraphX was used to digitalize the images. The white asterisks indicate the H cells showing the first GFP signal. The colored bar indicates the value of GFP fluorescence and cell volume. Scale bar, 50 μm. **C-F**, Ploidy distribution of *AT2G34910:H2A-GFP* positive cells in the 7-day-old (7 days after sowing) roots of WT (C) and *sim* (D), *ccs52a1* (E), and *sim ccs52a1* (F) mutants based on dual-color flow cytometry. **G-H,** Cross sections at the indicated distance of the root tip of 7-day-old roots of *SIM-GUS* (G) and *CCS52A1:CCS52A1-GUS* (H) seedlings. Scale bars, 20 μm.

### Both SIM and CCS52A1 control the ploidy level of root hair cells

SIM and CCS52A1 are pivotal regulators of endoreplication in Arabidopsis (Churchman et al., 2006; Vanstraelen et al., 2009). CCS52A1 contributes to root meristem maintenance, whereas the function of SIM in regulating root development, as well as the specific roles of SIM and CCS52A1 in root hair growth, remain elusive. To unravel the roles of SIM and CCS52A1 in Arabidopsis root endoreplication onset, we first delineated the expression patterns of *SIM* and *CCS52A1* using comparative cross sections from the *SIM:GUS* and *CCS52A1:CCS52A1-GUS* reporter lines. Confirming previous findings (Takahashi et al., 2013; Bhosale et al., 2018), initial expression of both genes could be observed in the BM (Fig. 2G and H). SIM accumulates first in the H cells, but in the TZ, its expression extended to the neighboring N and C cells. CCS52A1 was observed in both H and N cells, and C cells. For both genes, expression decreased in the EZ. The observed expression patterns of *SIM* and *CCS52A1* in the H cells suggested a potential involvement of both genes in the growth or endoreplication of H cells.

Primary root phenotypes of WT and mutants were compared to assess the functions of SIM and CCS52A1 during growth. Contrary to previous findings (Vanstraelen et al., 2009), no significant change in the root meristem length and size for the *ccs52a1* mutant compared to WT was observed under our growth conditions (Supplementary Fig. S4A-B), probably linked to the use of sugar-free rather than sugar-containing medium. Similarly, for the *sim* single and *sim ccs52a2* double mutants, no effects on meristem size and length were observed, as well as no observable differences in root width for all genotypes (Supplementary Fig. S4C). Accordingly, roots of the different mutants exhibited no significant changes in length compared to the WT during root growth, except for a slightly, but significantly, shorter *sim ccs52a1* root 7 days after germination (DAG) (Supplementary Fig. S4D). These observations indicate that SIM and CCS52A1 do not affect the overall root growth.

To analyze the effects of SIM and CCS52A1 on endoreplication in H and N cells, we mapped the DNA ploidy levels of both files in *sim* and *ccs52a1* single mutants, and *sim ccs52a1* double mutants (Fig. 2A and Supplementary Fig. S2). No outspoken differences could be observed between WT and mutants N cells, except for a 2-cell delay in the 2C-to-4C transition in the double mutant (Supplementary Fig. S2). For H cells, *sim* and *ccs52a1* single mutants lacked the accumulation of 16C cells in the EZ (Fig. 2A), and thus H cells remained at the 8C stage. The *sim ccs52a1* double mutant’s ploidy distribution resembled the patterns in these single mutants but initiated the 2C to 4C transition slightly later. Dual-color flow cytometry on mutants holding the *AT2G34910:H2A-GFP* marker line illustrated a delay in the 8C-to-16C transition in the *sim* primary root H cells, while *ccs52a1* and *sim ccs52a1* H cells remained at the 8C stage (Fig. 2D-F).

To investigate the effects of SIM and CCS52A1 on epidermal cell growth, MGX was used to map the cell volume in H and N cells of primary roots in the *sim*, *ccs52a1*, and *sim ccs52a1* mutants (Fig. 3, Supplementary Fig. S5). Mapping of N cell volume within the single mutants along the root axis revealed no major changes in maximum obtained cell volumes (Supplementary Fig. S5A). Only within the EZ, the average cell volumes appeared higher and lower than that of WT in *sim* and *ccs52a1*, respectively; the volume of *sim ccs52a1* N cells was significantly lower than that of the WT in both the TZ and EZ (Supplementary Fig. S5B and C). Differently, within the H cells, a reduction in cell volume was observed for all mutants in both the TZ and EZ, being most outspoken in the double mutant (Fig. 3). To elucidate the geometric factors leading to the decrease in cell volume, we mapped the cell length and width of WT and mutants H cells in both zones (Supplementary Fig. S6). The data showed that the mutation in *SIM* mainly affected the width of H cells, whereas the mutation in *CCS52A1* predominantly reduced H cell length. The *sim ccs52a1* double mutant displayed only a significant reduction in H cell length compared to WT.

**Fig. 3.**
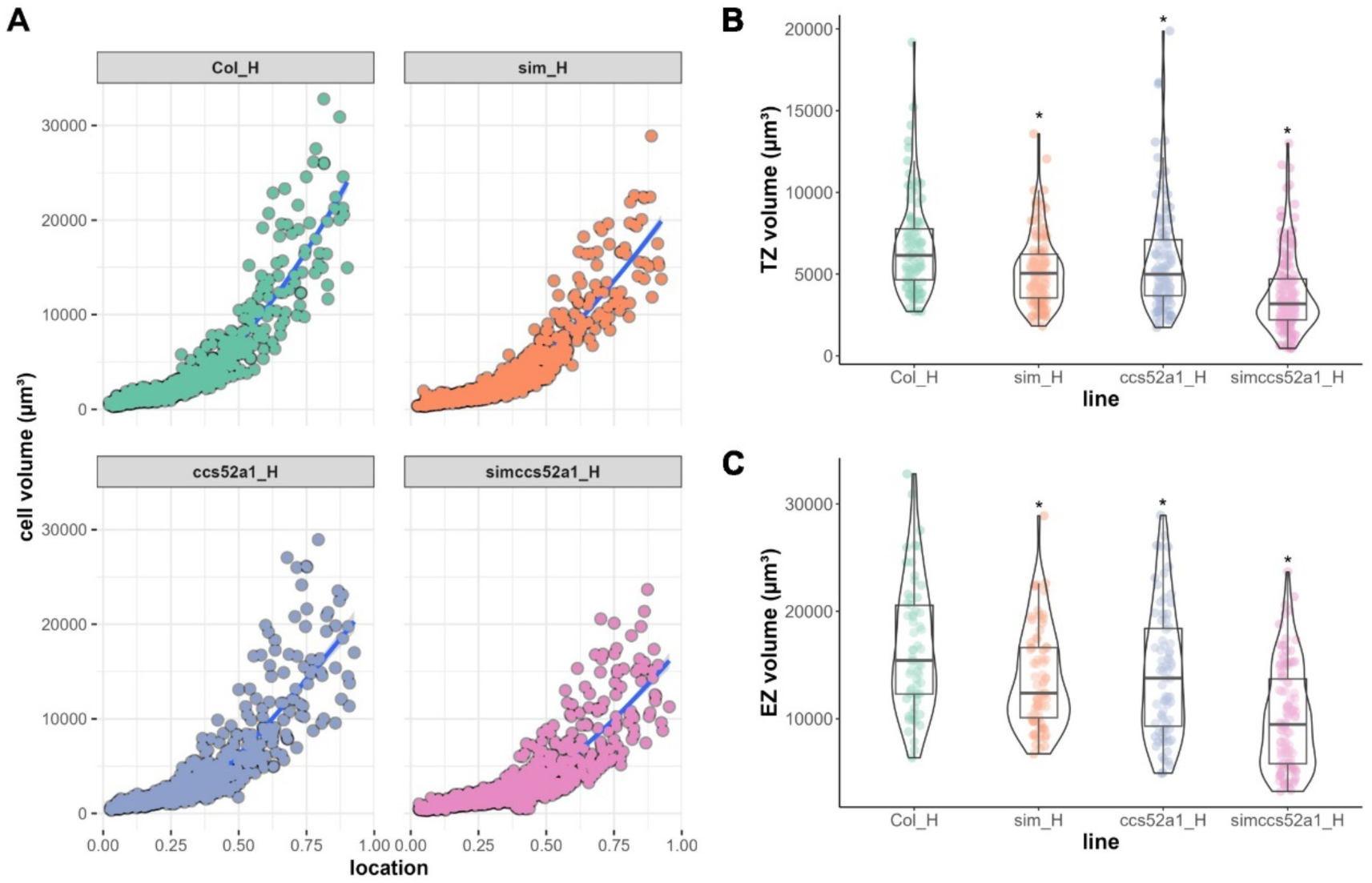
*SIM* and *CCS52A1* determine H cell volume in the primary root. **A**, Quantification and mapping of the H cell volume along the root longitudinal axis from QC (0.0 point) to the fifth/sixth elongated cell (1.0 point) in the primary roots of WT and *sim*, *ccs52a1*, and *sim ccs52a1* mutants. Blue lines show Loess regression curves. Over 600 cells among 3 independent roots for each genotype were measured. **B-C**, H cell volume of WT and mutants in the TZ (B) and EZ (C). The asterisks indicate a significant difference (adjusted p < 0.05) determined by nonparametric one-way ANOVA followed by Dunnett’s test post hoc test, compared to the WT (n ≥ 60 cells per genotype).

### SIM and CCS52A1 regulate cell expansion during root hair tip growth

Following entry in the EZ, tip growth of the H cell is initiated. To investigate the effects of SIM and CCS52A1 on this process, we first examined the endoreplication levels of the WT and mutants through the *VSR5:H2A-GFP* and *AT3G09330:H2A-GFP* reporters that are specifically expressed in the H cells from the EZ to the differentiated zone (Supplementary Fig. S3) and during the tubular-shaped hair formation (Supplementary Video S1), respectively. The expression patterns of these markers were unaffected in the *sim*, *ccs52a1*, and *sim ccs52a1* mutants compared to the control (data not shown). Dual-color flow cytometry with *VSR5:H2A-GFP* showed that the H cells from the 7-day-old WT root accumulated a ploidy level of 8C and 16C (Fig. 4A), whereas the *sim* and *ccs52a1* mutants displayed reduced H cell ploidy levels, with an increased portion of 4C cells (Fig. 4B-C). The *sim ccs52a1* double mutant exhibited an enhanced endoreplication defect, with the 16C population being barely detectable (Fig. 4D). Flow cytometric analysis using the *AT3G09330:H2A-GFP* reporter line confirmed that the 16C portion was reduced in *sim* and *ccs52a1*, and that H cells of the double mutant remained at the 8C stage (Fig. 4E-H). These findings demonstrate that SIM and CCS52A1 redundantly control the 8C-to-16C transition of maturing H cells.

**Fig. 4.**
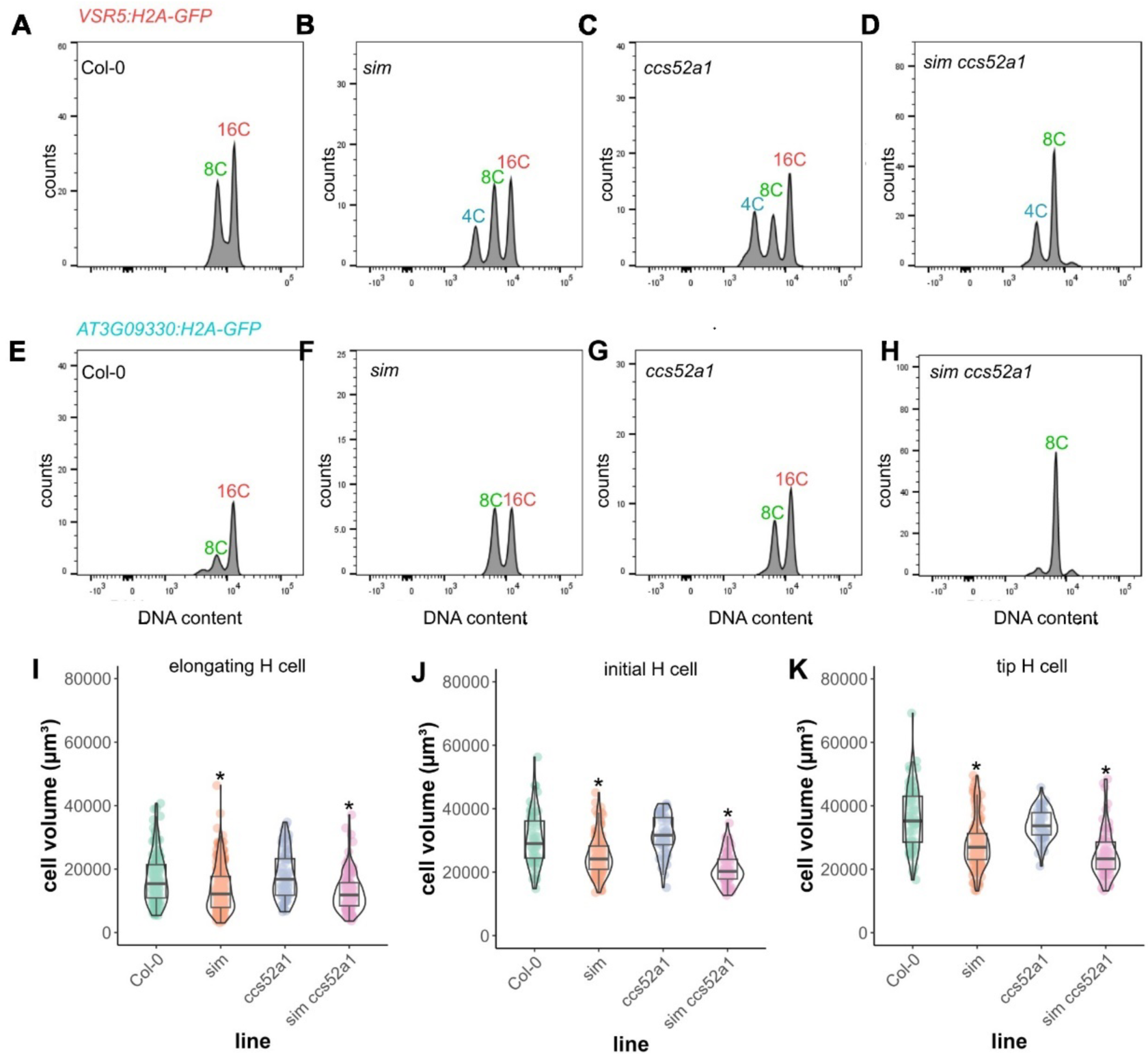
Impact of *SIM* and *CCS52A1* mutation on the ploidy level and volume of tip growing H cells. **A-H**, Ploidy distribution of *VSR5:H2A-GFP* (A-D) and *AT3G09330:H2A-GFP* (E-H) positive cells in 7-day-old roots of WT (A, E) and *sim* (B, F), *ccs52a1* (C, G) and *sim ccs52a1* (D, H) mutants based on dual-color flow cytometry. **I-K**, Volume of elongating (I), initial (J), and tip (K) H cells of WT and *sim*, *ccs52a1*, and *sim ccs52a1* mutants. Measurements were conducted on 5 independent roots per genotype (n ≥ 60 cells per genotype). Asterisks indicate a significant difference (adjusted p < 0.05) determined by nonparametric one-way ANOVA followed by Dunnett’s post hoc test, compared to the WT.

To correlate these findings with the H cell volumes, we performed MGX on WT and mutant roots beyond the EZ, including cells just prior to tip growth, at the tip growth initiation stage, and when tip growth was completed. In the *sim* and *sim ccs52a1* mutants, the volumes of all H cells were significantly reduced compared to the WT, while no significant changes were observed in *ccs52a1* (Fig. 4I-K). These findings indicate that primarily *SIM* contributes to the H cell volume in the differentiated root zone.

To assess the effects of *SIM* and *CCS52A1* on root hair tip growth, root hair parameters were measured in the WT and mutant lines (Fig. 5A-D). Within the *ccs52a1* mutant, the first visual tip grown cell was found more distantly from the root tip compared to the WT (Fig. 5E), indicative for a delayed root tip cell differentiation. In all mutants, the final root hair length was significantly reduced compared to the WT (Fig. 5F). In addition, time-lapse imaging showed that the root hair growth rate was significantly decreased in the *ccs52a1* and *sim ccs52a1* mutants compared to both WT and *sim* (Fig. 5G), indicating a positive effect of *CCS52A1* on the root hair growth rate.

**Fig. 5.**
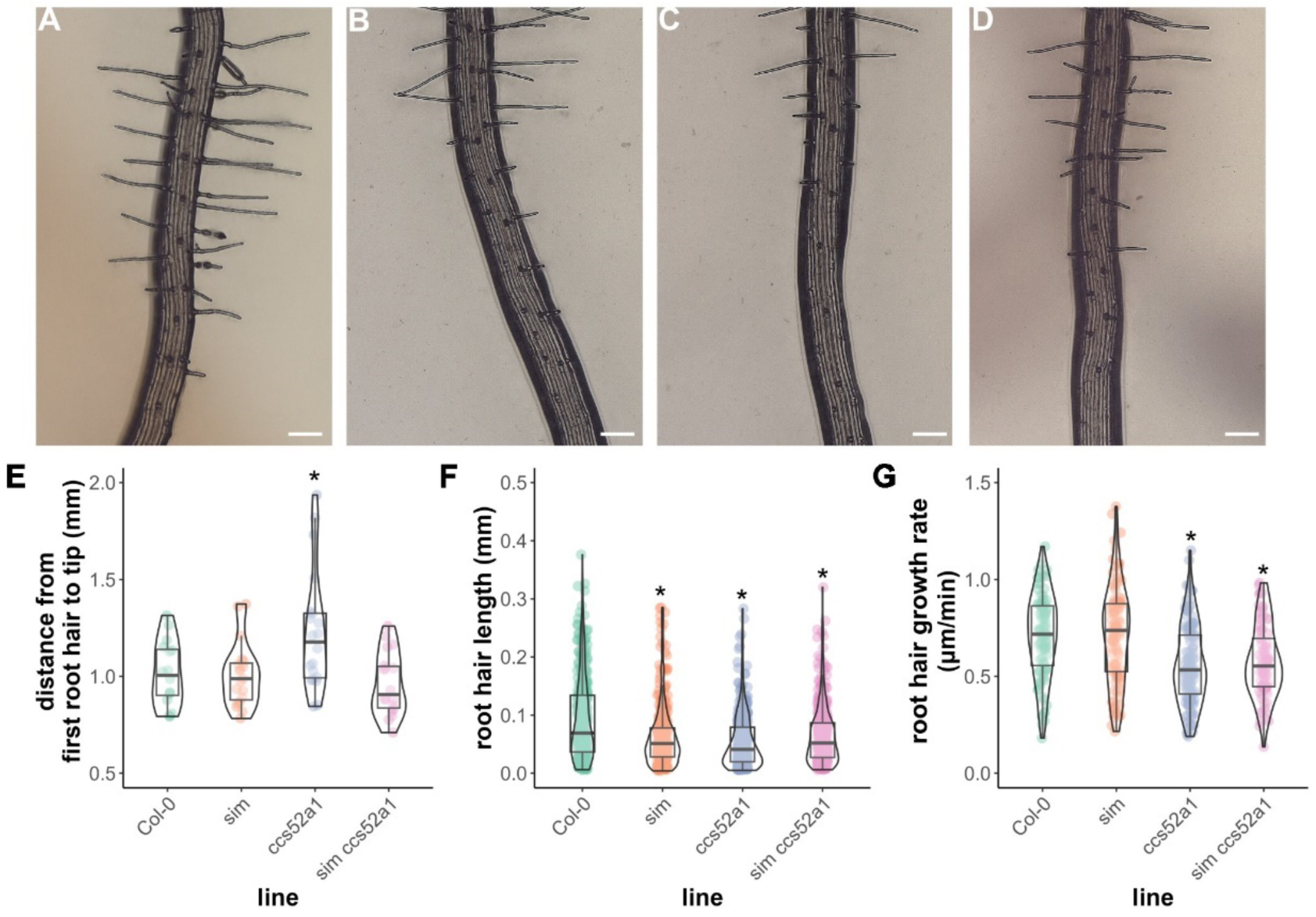
*SIM* and *CCS52A1* control root hair tip growth. **A-D**, Root images used for phenotype quantification from WT (A), *sim* (B), *ccs52a1* (C), and *sim ccs52a1* (D). Scale bars, 0.1 mm. **E-G**, Quantitative measurements of the distance from the root tip to the first visible root hair (E), root hair cell length (F), and root hair growth rate (G). Data were obtained from over 15 independent roots per genotype (n ≥ 80 cells per genotype). Asterisks indicate significant difference (adjusted p < 0.05) between WT and mutants. For root hair cell length and growth rate, the statistical analysis was done by nonparametric one-way ANOVA, followed by Dunnett’s post hoc test; for distance from the first root hair to tip, the statistical analysis was done by parametric one-way ANOVA, followed by Dunnett’s post hoc test.

### Cell wall composition changes in the primary roots of endocycle mutants

During cell expansion, the plant cell wall is subjected to modifications. In light of the observed decrease in the volume of H cells in the primary roots of the *sim* and *ccs52a1* mutants (Fig. 3), we investigated potential modifications in the primary root cell wall, induced by *SIM* and *CCS52A1* mutation by employing the comprehensive microarray polymer profiling (CoMPP) method (Moller et al., 2012; Rydahl et al., 2018). At seven days after sowing, primary roots from WT and *sim*, *ccs52a1*, and *sim ccs52a1* mutants were collected, cell walls were extracted and fractioned into a pectin-rich and hemicellulose-rich (fraction using 1,2-cyclohexylenedinitrilotetraacetic acid (CDTA) and NaOH, respectively). The two fractions were printed as microarrays and probed with monoclonal antibodies (mAbs) that target specific cell wall polysaccharide and glycoprotein epitopes. By quantifying the intensity of each microarray spot, this method allowed to semi-quantify the cell wall composition difference between the WT and the different mutants (Table 1).

**Table 1.**
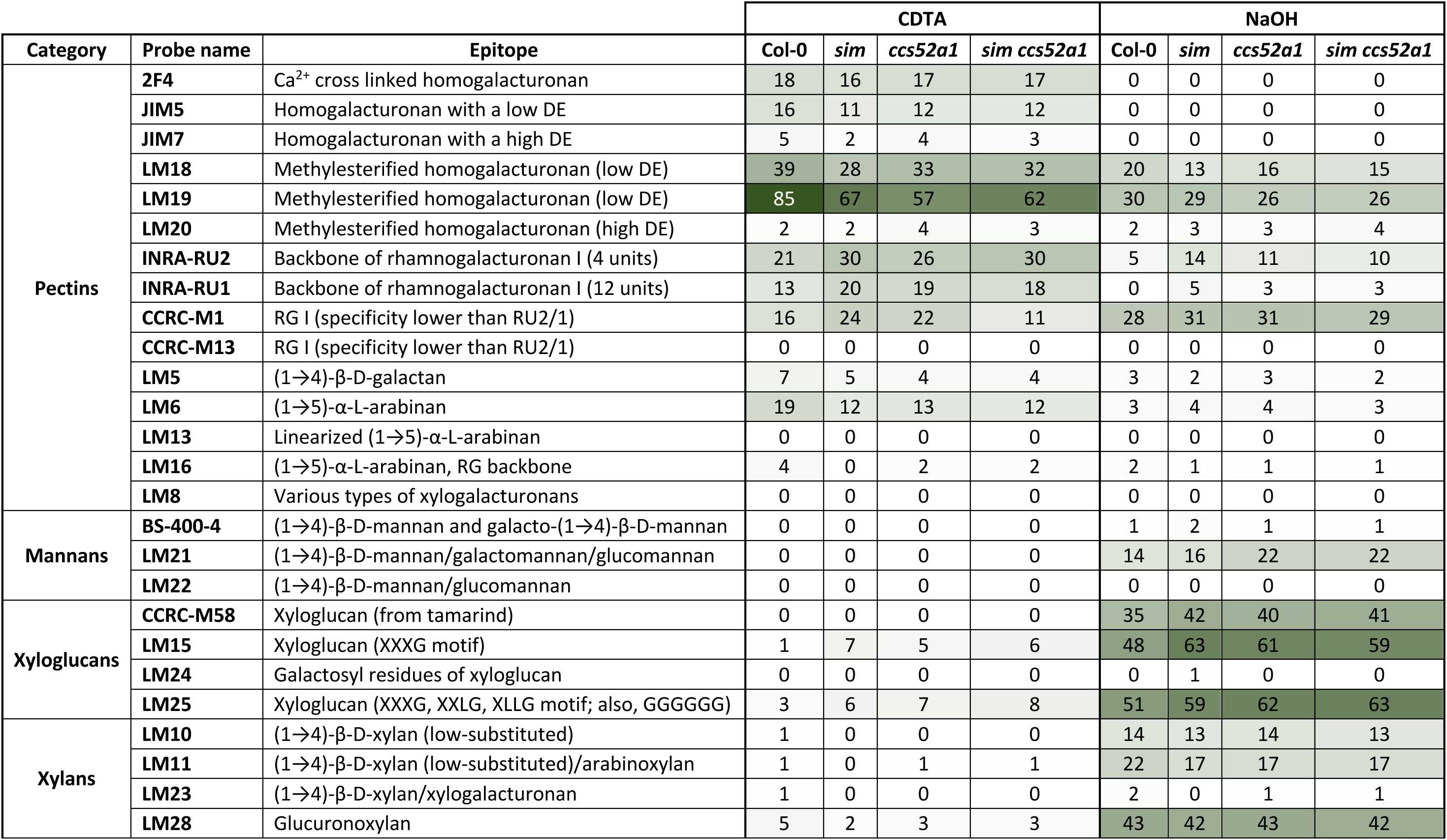
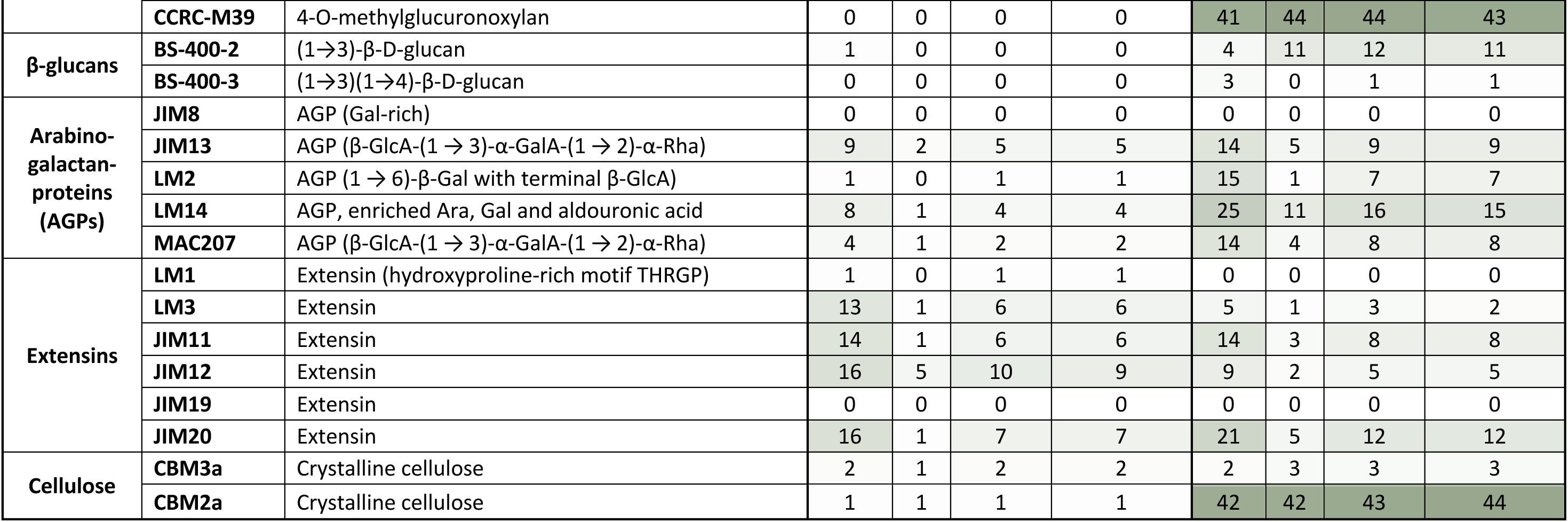
Carbohydrate microarray on different extraction of roots of WT and mutants. Pectin and hemicellulose were extracted using 1,2-cyclohexylenedinitrilotetraacetic acid (CDTA) and NaOH, respectively. The spot value represents the signal of intensity of each immune antibody, with the higher intensity of green coloration indicating higher values. All spot intensity values are normalized to the highest fluorescence value. n = 6 pools of 2 biological repeats. DE, degree of methylesterification.

Statistical analysis showed a significant reduction of HG with a low degree of methylesterification (DE) (recognized by probes LM18 and LM19), (1→5)-α-L-arabinan (LM6), and extensins (JIM12, JIM20, and LM3) in the *sim*, *ccs52a1*, and *sim ccs52a1* mutants compared to the WT (Fig. 6A-F, Table 1). The DE value of HG is crucial in determining plant cell expansion (Pelletier et al., 2010; Haas et al., 2020). Cell wall-resident PMEs demethylate HG during the process of cell expansion, resulting in a decrease in the DE value of HG (Pelletier et al., 2010; Kohorn, 2016). To compare the demethylesterification levels of HG in the root hairs between WT and tested mutants, we conducted immunolocalization staining using the LM19 antibody (Fig. 6G-N). Compared to the WT, the *sim*, *ccs52a1*, and *sim ccs52a1* mutants experienced a reduction of demethylesterified HG in the root hairs (Fig. 6O), suggesting that SIM-and CCS52A1-mediated endoreplication may promote root hair elongation by affecting the demethylesterification pattern of HG.

**Fig. 6.**
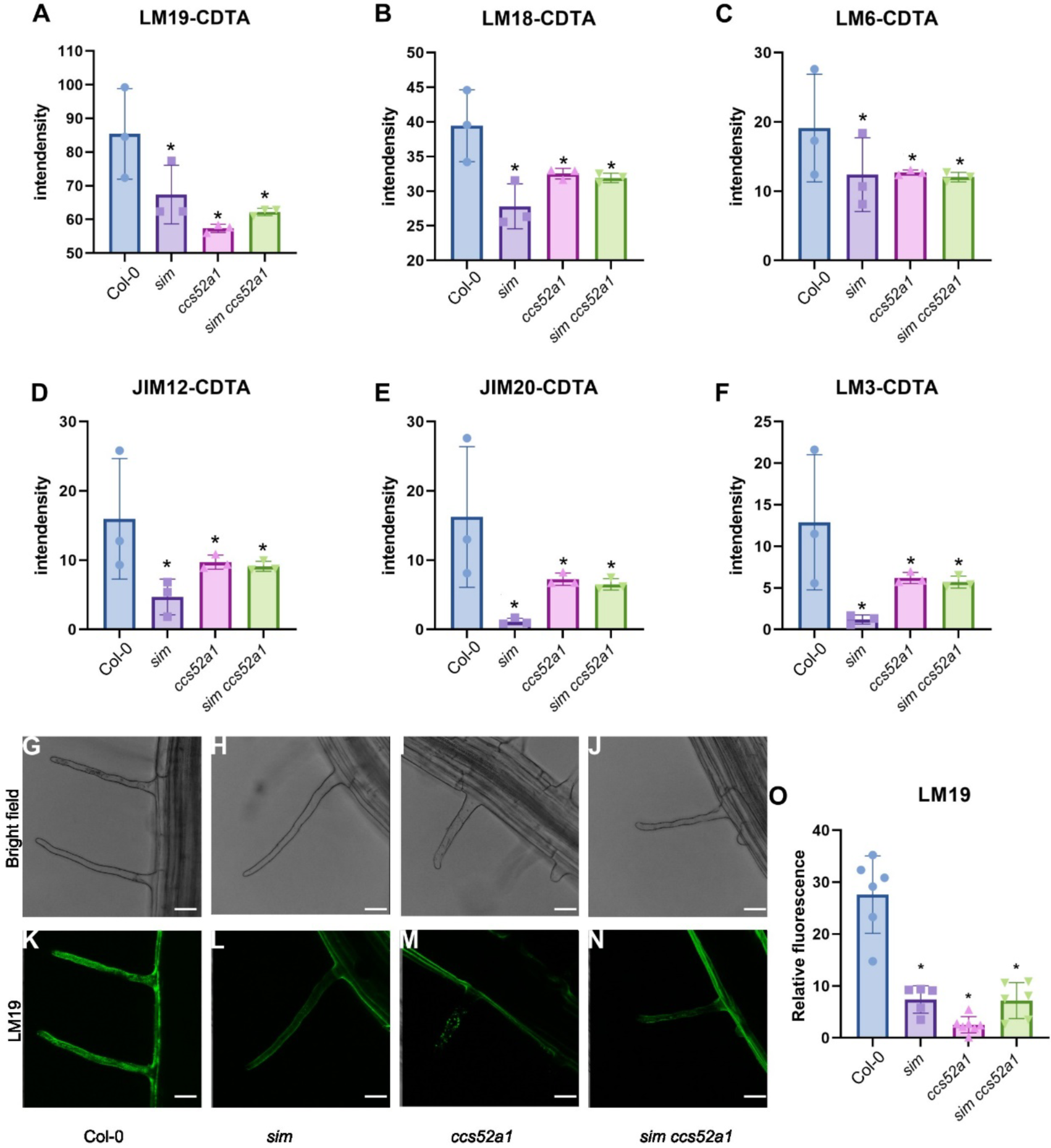
Impact of *SIM* and *CCS52A1* mutation on primary root cell wall modifications. **A-F**, Quantification of LM19 (A), LM18 (B), LM6 (C), JIM12 (D), JIM20 (E), and LM3 (F) intensity signal in cell wall extracts of WT (Col-0), *sim*, *ccs52a1*, and *sim ccs52a1* 7-day-old roots. Asterisks denote significant differences compared to the control (p < 0.05) determined by two-way ANOVA followed by Dunnett’s post-hoc test. **G-N**, Bright-field (G-J) and fluorescence (K-N) images of root hairs of WT and *sim*, *ccs52a1*, and *sim ccs52a1* mutants stained with the monoclonal antibody LM19 (green fluorescence). Scale bars, 20 µm. **O**, Quantification of the LM19 fluorescence intensity. Asterisks indicate statistical significance (adjusted p < 0.05) compared to WT, determined by parametric one-way ANOVA and Tukey’s post-hoc test (n > 3 roots per genotype).

## Discussion

The relationship between endoreplication and H cell growth is still unclear. DNA topoisomerase VI complex mutants, which fail to progress beyond the 8C ploidy level, were found to display a disturbed root hair tip growth (Sugimoto-Shirasu et al., 2005; Kirik et al., 2007). Conversely, using a diverse set of H cell mutants, Sliwinska et al. (2015) found no clear correlation between root hair length and the level of endoreplication. The results of the latter study may be potentially hampered by the fact that mutants were selected based on root hair phenotypes, with likely other processes than the endocycle being the primary cause of the root hair phenotype. Therefore, studying mutants in genes specifically controlling endoreplication may give a more conclusive answer to the question about the relationship between H cell growth and ploidy level. Here, using three such mutants, we mapped the DNA ploidy level along the complete root hair cell file using two independent methods, being quantification of DAPI-stained nuclei and dual-color flow cytometric analysis. Subsequently, these data were combined with cell volume measurements obtained through MGX, enabling the exploration of the effect of endoreplication on the H cell phenotype.

### SIM and CCS52A1 regulate the H cell volume

Both ploidy measurement methods consistently demonstrated that WT H cells enter the 8C and 16C stages after they leave the meristematic zone. This finding aligns with the results reported by Pasternak et al. (2022) and our previous transcriptome-based modeling of DNA ploidy levels along the complete root (Bhosale et al., 2018). We also confirmed that H cells undergo an additional round of endoreplication compared to their neighboring N cells. Notably, not only do H cells obtain their final ploidy level before root tip growth, as illustrated for collet hairs (Sliwinska et al., 2015)), they also were found to hold a 4C DNA content within the BM preceding a rapid increase in volume, suggesting that endocycle onset may be a prerequisite for rapid H cell elongation from the moment they exit the cell cycle. This hypothesis was tested through phenotypic analysis of *sim* and *ccs52a1* mutant root hairs.

Arabidopsis SIM and CCS52A1 have previously been recognized as pivotal regulators of endoreplication in trichomes, roots, sepals, and leaves (Churchman et al., 2006; Vanstraelen et al., 2009; Robinson et al., 2018). Within the root, consecutive cross section images revealed that *SIM* and *CCS52A1* are expressed in H cells from the BM onwards, indicating that both genes may regulate endocycle onset and progression within the H cells. Indeed, both single mutants displayed a clear reduction in the number of cells holding a 16C ploidy content, compared to WT plants. It suggests that SIM and CCS52A2 independently control the 8C-to-16C transition, probably through targeting endocycle-required CDK and cyclin subunits, respectively. No clear effect on the 4C-to-8C transition was observed, whereas the 2C-to-4C transition was only slightly delayed in the *sim ccs52a1* double mutant. These data suggest the presence of other H cell endocycle modulators. Likewise, besides the 2-cell delay in the 2C-to-4C transition in the double mutant, there was no clear effect on the 8C-to-16C transition in the mutant N cells, suggesting again the existence of other tissue-specific endocycle regulators. Putative candidates can be found among other members of the large SIM/SMR gene family, including *SMR1*, *SMR2*, *SMR9*, and *SMR13*, for which mutants have been demonstrated to control the primary root meristem size in a cooperative manner (Nomoto et al., 2022; Goldy et al., 2023). A key candidate gene for the endocycle control in N cells is *SMR1*, showing a more prominent expression in N compared to H cells (Bhosale et al., 2018).

To investigate the effects of SIM and CCS52A1 on H cell expansion, we mapped the H cell volume along the root longitudinal direction. Both mutants showed a reduction in cell volume, being most outspoken in the EZ, hence the region with a clear quantitative effect on the endocycle and thus indicative for a supportive role for the endocycle in the cellular growth. Interestingly, geometrical analysis showed that SIM and CCS52A1 promote cell expansion in the transversal and longitudinal directions, respectively. Therefore, SIM and CCS52A1 may modulate H cell expansion in different directions in a yet to be determined way, both resulting in a change in cell volume. Anisotropic cell expansion is commonly attributed to cell wall changes, including the pectin demethylation pattern and organization of cellulose fibers (Peaucelle et al., 2015; Bou Daher et al., 2018), the latter being determined by the trajectory of cellulose synthase complexes imposed by cortical microtubuli. Within meiotic and mitotically dividing cells, different CDKs and cyclins have been localized to microtubule arrays and demonstrated to impact their dynamics (Weingartner et al., 2003; Sofroni et al., 2020). Consequently, it is possible that in post-mitotic cells, CDK/cyclin complexes, which could be targeted by SIM or CCS52A1, likely govern the direction of H cell expansion. This regulation could occur by modulating the function of microtubules, likely through the phosphorylation of microtubule-associated proteins that control microtubule dynamics. Interestingly, the potential APC/C substrate PIKMIN1 was found to localize at cortical microtubuli (Willems et al., 2023).

In addition to a reduction in H cell volume in the TZ, root hair length of all mutants was slightly reduced, confirming the shorter mature hair length phenotype of the *ccs52a1* mutant reported before (Takahashi et al., 2013). These data suggest that obtaining a 16C DNA may be essential to achieve both a WT cell volume and root hair length. Interestingly, time-lapse recording showed that *ccs52a1* and *sim ccs52a1* mutants experienced a decreased root hair growth rate compared to the control. Recently the *CDK-like 12 (CDKL12)/BUZZ* gene was identified as an evolutionary conserved regulator of root tip growth (Lehman et al., 2023). Transcriptome analysis suggests that *CDKL12/BUZZ* may operate through control of RNA biogenesis and ribosome functioning, as well as in actine and tubulin dynamics, and cell wall dynamics, all processes that have been linked to endoreplication (Bhosale et al., 2019). Thus, it will be of interest to test whether CDKL12/BUZZ or its interacting cyclins are targeted by SIM or CCS52A1.

Within the EZ, we also observed alterations in the N cell volume for *sim*, *ccs52a1*, and *sim ccs52a1* mutants compared to the WT. The N cell volume in the *sim* mutant was found to be greater than that of WT, while the N cell volumes in both *ccs52a1* and *sim ccs52a1* mutants were diminished. As no outspoken ploidy changes were observed, these changes must have arisen in an endocycle-independent manner. Probably, the N cell volume changes arise as a reaction on ploidy-driven cell geometrical changes of the neighboring H cell. The mutation in *SIM* led to a narrowing of the H cell width compared to WT, potentially providing the adjacent N cells with more room for transversal expansion. The H cell width in *ccs52a1* and *sim ccs52a1* mutants remained unaffected, indicating that *CCS52A1* may play a distinct role in the neighboring N cells. Despite these observations, the current evidence is insufficient to confirm these hypotheses. We propose that integrating computational studies with live-imaging techniques focused on cell volume could offer a more robust approach to address these hypotheses.

### SIM and CCS52A1 modulate the root hair tip growth

An argument against control of tip growth by endoreplication through manipulation of microfilaments is the lack in the *ccs52a1* and *sim* mutants of aberrant hair structures frequently seen in actin mutants (Wan et al., 2017; Chin et al., 2021). Alternatively, recent studies have identified modifications in the shank-localized cell wall during root hair elongation (Herburger et al., 2022; Schoenaers et al., 2024). The genome content can affect cell wall composition, consequently influencing cell volume dynamics (Narukawa et al., 2015; Corneillie et al., 2019). For example, a mutation in the *KAKTUS* (*KAK*) gene promoted endoreplication in the vascular cells, accompanied by modifications in cell wall *O*-acetylation (Bensussan et al., 2015). Likewise, a recent study demonstrated that reduced endoploidy levels in the *ccs52a2* mutant lead to a reduction in the methylesterification level of pectin in the hypocotyl hook, accompanied by a decrease in cell expansion (Ma et al., 2022). Pectin is a linear polymer and its major domain HG is composed of α-1,4-linked-D-galacturonic acid (Saffer, 2018). In plant cells, pectin is initially synthesized in the Golgi complex in a highly methylesterified form. During cell expansion, PMEs remove methyl groups, causing a reduction in methylesterification of pectin with negative charges (Kohorn, 2016). Increasing evidence revealed that demethylated pectin plays a positive role in regulating plant cell expansion (Pelletier et al., 2010; Cheong et al., 2019; Haas et al., 2020). One possible explanation is that the decrease in methylesterification of pectin reduces the cell wall pH, which in turn promotes the activity of expansins (Cosgrove, 2024). Our study reveals that the mutations in *SIM* and *CCS52A1* reduce the signal of demethylated HG in primary roots, which may explain the observed decrease in H volume in the primary roots of the *sim*, *ccs52a1*, and *sim ccs52a1* mutants compared to that of WT. In addition, further alterations of the cell wall composition, including (1→5)-α-L-arabinan and extensins, could also be attributed to the altered cell expansion pattern. For example, arabinans are considered to act as cell wall ‘plasticizers’ (Moore et al., 2013) and their lower abundance in the mutant plants studied might help to explain the reduced cellular expansion, leading to a decrease in cell volumes. Extensins, which exhibited lower detection signals in mutant plants, are important for cell wall assembly as they interact with pectin. For example, EXT3 and pectin form coacervates that serve as a template for the deposition of further cell wall material (Cannon et al., 2008). Finally, all mutants showed higher signals for xyloglucan. Xyloglucan is a major hemicellulose in cell walls of non-grass vascular plants and consists of a β-(1→4)-linked glucose backbone decorated with short xylose-rich branches. Xyloglucan knockout mutants show a decreased cell wall creep under growth-inducing conditions and strong root hair phenotypes (bulging, reduced growth rates, and length), but plants still grow appreciably (Park and Cosgrove, 2012), likely assisted by compensation mechanism such as upregulated pectin production (Sowinski et al., 2022). Analogously, it is tempting to speculate that endoreplication-induced changes in the pectin methylation pattern are compensated by an increased xyloglucan content, which would help explaining the absence of strong root hair phenotypes in *sim*, *ccs52a1*, and *sim ccs52a1* mutants. These findings suggest a link between H cell expansion regulation by SIM and CCS52A1 and cell wall modifications.

## Conclusion

The effect of endoreplication on cell volume has long been debated. In our study, we found that Arabidopsis root H cells display a close relationship between cell volume and ploidy. H cells’ 4C and 16C ploidy increases preceded the rapid cell volume increase in the EZ and the initiation of root hair tip growth, respectively. Moreover, both *SIM* or *CCS52A1* loss-of-function decreased the volume of TZ and EZ cells, which was enhanced in the double mutant. With both genes playing a likely specific role in ploidy control, such direct consequences of *SIM* or *CCS52A1* mutation on cell volume are strongly supportive for a contribution of endoreplication to cell growth. However, even in the double mutant cells lacked a highly outspoken decrease in cell volume phenotype. This might be in part explained by the fact that cells still reached an 8C DNA content. This suggests that, in addition to SIM and CCS51A1, additional players are involved in controlling the ploidy level of H cells, and effects on cell volume might be even more pronounced in a mutant completely deficient in endocycle onset. The mechanism by which SIM and CCS52A1 regulate H cell expansion during endoreplication remains unclear, but accumulating evidence links an increase in endoploidy to cell wall modifications (Bhosale et al., 2019; Ma et al., 2022; Goldy et al., 2023) that, through an interplay with turgor-driven vacuole expansion, may influence final cell size. The mechanistic framework for how a change in ploidy results in such a specific cell wall modification remains an open question.

## Materials and methods

### Plant material, growth conditions and chemical treatments

*Arabidopsis thaliana* lines (Supplementary Table S1) used are in the Columbia-0 (Col-0) background. Sterilized seeds were plated on a sugar-free medium consisting of 1/2 Murashige and Skoog (MS) medium supplemented with 1% plant agar and 0.5 g/L MES, adjusted to pH 5.8. The seeds were stratified in the dark for three days at 4°C and subsequently transferred to a growth room with a 16-h light/8-h dark cycle (70 μmol m^-2^s^-1^) at 22°C for seven days.

The transgenic lines (*AT2G34910:H2A-GFP*, *VSR5:H2A-GFP*, and *AT3G09330:H2A-GFP*) expressing the H2A-GFP fusion protein specifically in the Arabidopsis root H cells were constructed with the Golden Gate cloning with the GreenGate method. Six entry modules [pGG A-promoter-B, pGG B-linker-C, pGGC-H2A (no stop codon)-D, pGGD-GFP-E, and pGGE-G7 terminal-F] were assembled into a destination (pFASTR-A-G) with the restriction enzyme BsaI and T4 DNA ligases. In summary, the A-B entry module contains the tissue-specific promoter, the C-D module contains histone A without stop codes, and the D-E module contains the GFP fluorescence tag, and the E-F module contains the terminal. The detailed method is described by Decaestecker et al. (2019). The promoter sequences in the pGG A-promoter-B module are described in Supplementary Table S2.

### Confocal microscopy

High-resolution images were obtained using an LSM 710 confocal laser-scanning microscope (Zeiss) equipped with a ×40 water immersion objective lens (NA 0.8). Cell outline staining and detection settings adhered to the ClearSee protocol (Kurihara et al., 2015). Fluorescence emission of the root hair marker lines was detected at 575 nm for PI and with a 500-nm and 530-nm bandpass filter for eGFP. eGFP was excited at 488 nm, and emission was detected using a 500-nm and 530-nm bandpass filter. Live imaging of *AT3G09330:H2A-GFP* Col-0 (Supplementary Video S1) was performed using an optical chamber and an LSM 900 confocal laser-scanning microscope (Zeiss) equipped with a ×20 air objective lens (NA 0.8).

### 3D analysis by MGX

The acquired images were converted to “tif” format using Fiji software for subsequent 3D segmentation with MorphoGraphX (version 2.0) (Montenegro-Johnson et al., 2015; Wolny et al., 2020). To enhance image quality, the images were smoothed using a Gaussian blur (x=0.6, y=0.6, z=0.3) before being segmented using ITK=1000. Manual corrections were made to address segmentation mistakes. The 3D cellular mesh was created with a cube size of 1.0.

Geometrical parameters, including cell volume, length, and width, were analyzed using the “heatmap” function. For position analysis of the roots of the WT and the *sim*, *ccs52a1*, and *sim ccs52a1* mutants, the “3DCellAtlas” function of MGX (Strauss et al., 2022) was applied. This involved the creation of a 2.5D surface mesh. Mapping analysis was performed using the “ggstatsplot” R package, with the 0.0 point set at the QC and the 1.0 point set at the fifth/sixth elongated cells.

### Endoploidy mapping analysis

The nuclei of 7-day-old roots were stained with DAPI solution, while the cell outlines were stained using SR2200 (Musielak et al., 2016; Tofanelli et al., 2019). The DAPI-stained nuclei were imaged in Z stacks using a LSM-710 confocal microscope (Zeiss). Nuclear sizes were extrapolated from images of the center of targeted nuclei. The average number of pixels of QC nuclei served as the 2C reference. DAPI-stained nuclei were collected from around 10 H and N files from 4 to 5 independent seedlings for each genotype.

### Flow cytometer experiment

The endoploidy levels of WT and mutant (*sim*, *ccs52a1*, and *sim ccs52a1*) H cells were measured in 7-day-old Arabidopsis roots using flow cytometric analysis of H cell-specific marker lines with a H2A-GFP tag. Multiple marker lines cover different H-cell developmental stages. For flow cytometry on nuclear GFP lines, cut root tips were chopped with a razor blade in 1 mL Galbraith’s buffer (45 mM MgCl_2_, 20 mM MOPS, 30 mM sodium citrate, 0.1% Triton X-100, pH = 7.0 with NaOH), then filtered through a 30-µm CellTrics filter. The DNA was stained with 1 μL (1 mg/mL) DAPI solution in the filtered supernatants for flow cytometric.

Flow cytometry (Quantum Pro-Flow-Cytometer) was performed on distinct regions of each root hair marker line based on the marker expression pattern. DAPI-stained nuclei were excited by illumination at 405 nm and equipped with an additional 488-nm laser to excite and detect GFP-specific fluorescence. The flow cytometric results were analyzed by FlowJo software.

### Time-lapse recording of root hair growth

Seven-day-old plants growing in a sugar-free medium were positioned on the stage of the microscope, and time-lapse sequences capturing hair growth were recorded using NVX10. Hair growth measurements were conducted by tracing hair position at fixed time intervals (5 min) to record the root hair growth over a span of 5 h.

### Statistical analysis and quantification

Statistical analysis was performed using the “ggstatsplot” package in R. Samples size, number of biological replicates and statistical tests used are detailed in Supplemental Dataset S1. The significance threshold was determined at an adjusted p < 0.05, denoted by one asterisk. Multiple testing correction was executed following the Benjamini–Hochberg procedure with α = 0.05 (Fridman et al., 2021).

### Cross-section GUS staining

To investigate the expression pattern of *SIM* and *CCS52A1* in the primary root of WT, cross-section staining was performed on the reporter lines of these two genes 7 DAG, following the protocol described in Bhosale et al. (2018).

### Material preparation for comprehensive microarray polymer profiling (CoMPP)

The primary roots of both WT and mutants were harvested 7 DAG and immediately frozen in liquid nitrogen. The freeze-dried materials were homogenized in 2-mL Eppendorf tubes with a metal ball. The finely homogenized lysate was mixed with 70% ethanol, vortexed thoroughly, incubated for 30 min, and centrifuged at maximum speed for 10 min. This process was repeated three times. The pellet was collected and resuspended in acetone, followed by air-drying overnight, resulting in a whitish powder (=alcohol insoluble residue, AIR). The extraction weight was measured, with each sample weighing at least 5 mg.

### Extraction of cell wall glycans

To extract the pectin-rich fractions from the AIRs, 500 μl of 50 μM CDTA (pH 7.5) was added to each plant sample. After brief vortexing, samples were homogenized using a TissueLyser, followed by centrifugation for 1 min at the max speed. The supernatant was carefully transferred into a new Eppendorf tube and stored at 4 °C. The hemicellulose-rich fraction was sequentially extracted from the AIR by incubating the washed pallet with 500 μl of 4 M NaOH containing 0.1% w/v NaBH_4_. The supernatants were diluted in a 1:2, 1:20, and 1:50 concentration for the next step.

### Printing microarrays, probing, and quantification

Arrays were printed on nitrocellulose sheets using an ArrayJet Sprint microarray printer (ArrayJet, Roslin, UK; Rydahl et al., 2018). Arrays were blocked (5% skimmed milk powder in Tris-buffered saline with Tween 20 (TBST): 20 mM Tris-HCl, 140 mM NaCl, pH 7.5, 0.1% Tween 20) for 1 h, followed by incubation with primary cell wall antibodies or carbohydrate binding modules 1:100 in blocking solution for 1.5 h, followed by washing with TBST (3 x 5 min). Next, the secondary antibodies (BCIP/NBT; 1:1000, diluted with the blocking solution) were added and incubated for 1.5 h, followed by washing.

BCIP/NBT color development was quantified based on the gray integrated density of each colored precipitate point.

### Immuno-fluorescence labeling

Methylesterified HG (low DE) was detected using an LM19 mAb (PlantProbes). For in situ immuno-fluorescence, 7-day-old seedlings grown in channel slides (ibidi, 80601) were fixed in PBS containing 2.5% (v/v) glutaraldehyde. Channels were then thoroughly rinsed with 3 ml of PBS applied with a syringe under constant pressure for 30 s to remove GA. Seedlings were blocked in PBS with 2% (w/v) BSA for 30 min, rinsed with a syringe, and then incubated with LM19 (diluted 1:20 in PBS) at 22.5°C for 2 h. After rinsing, a secondary antibody (diluted 1:50 in PBS; Invitrogen, Alexa Fluor 488 goat anti-Rat IgM (Heavy chain) cross-adsorbed secondary antibody (A-21212)) was added and incubated for 60 min at room temperature. Samples were rinsed with a syringe and subjected to confocal microscopy (Leica SP5; laser line 488 nm, EM 520–530 nm). This protocol has followed the procedure outlined by Herburger et al. (2022).

## Acknowledgments

The authors thank Prof. Richard Smith on MorphoGraphX, Qian Zhao on data statical analysis, and Annick Bleys for critical reading and manuscript preparation.

## Author contributions

L. L. and L.D.V. designed the research; L.L. performed the experiments with technical assistance from I.V.; K.H. performed the immunolabelling work; I.V. performed the cross-section GUS analysis; L.L. analyzed the experimental data and wrote the manuscript; L.D.V. and J.H. supervised the writing.

## Supplementary data

The following materials are available in the online version of this article.

**Supplementary Fig. S1.** Geometrical analysis in WT primary root epidermal cells.

**Supplementary Fig. S2.** *SIM* and *CCS52A1* only marginally control the ploidy level of N epidermal cells.

**Supplementary Fig. S3.** Expression patterns of *AT2G34910:H2A-GFP* and *VSR5:H2A-GFP* marker lines in the root.

**Supplementary Fig. S4.** Primary root phenotyping of endocycle mutants.

**Supplementary Fig. S5.** Quantification of the effects of *SIM* and *CCS52A1* on N cell volume in the primary root.

**Supplementary Fig. S6.** Quantification of effects of *SIM* and *CCS52A1* on the length and width of H cells in the primary root tip.

**Table S1.** Arabidopsis lines used in this study.

**Table S2.** Sequences of promoters used in root hair-specific marker lines.

**Supplementary Video S1:** Expression of *AT3G09330:H2A-GFP* marker line.

**Supplementary Dataset S1:** Statistical analysis

