## Supplemental figure for "*SIM* and *CCS52A1* control root hair cell endoreplication and expansion"

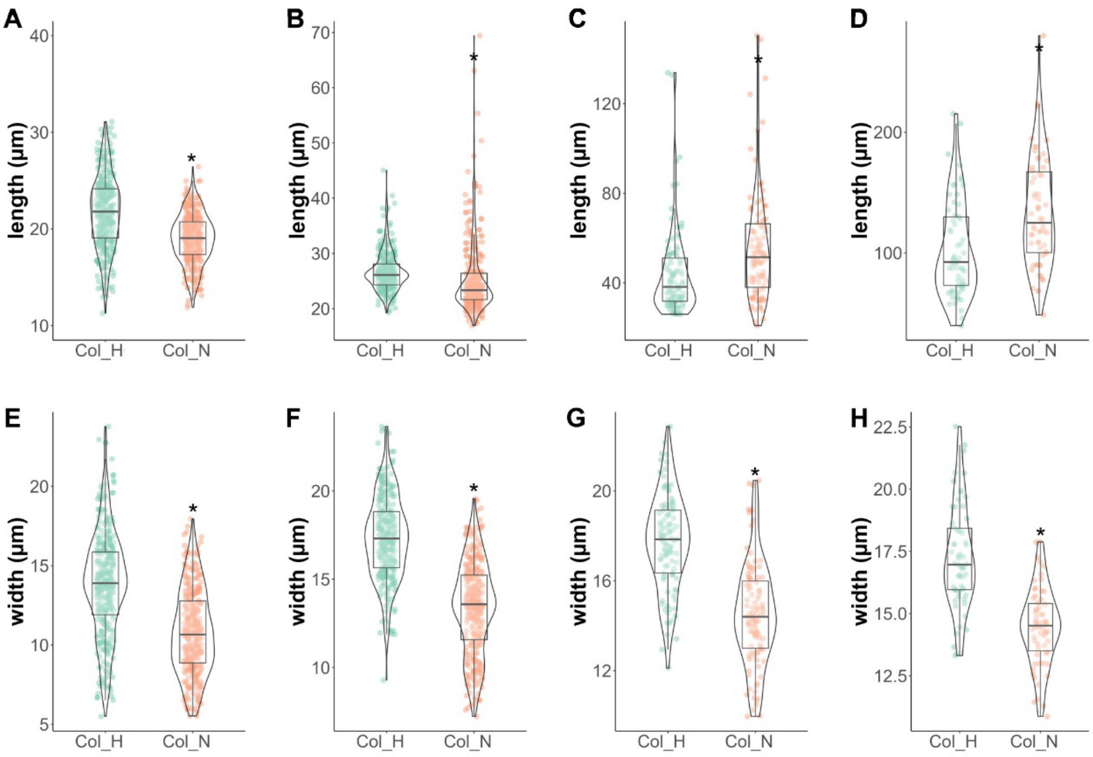


**Supplementary Figure S1. Geometrical analysis in WT primary root epidermal cells. A-H**, Cell length (A-D) and width (E-H) of H and N cells within the proximal (A, E) and basal (B, F) meristem, and transition (C, G) and elongation (D, H) zone. Over 600 cells from 3 independent roots for each cell type were measured. Asterisks denote significant differences (adjusted $p < 0.05$) determined by parametric t-test followed by Tukey's post hoc test.

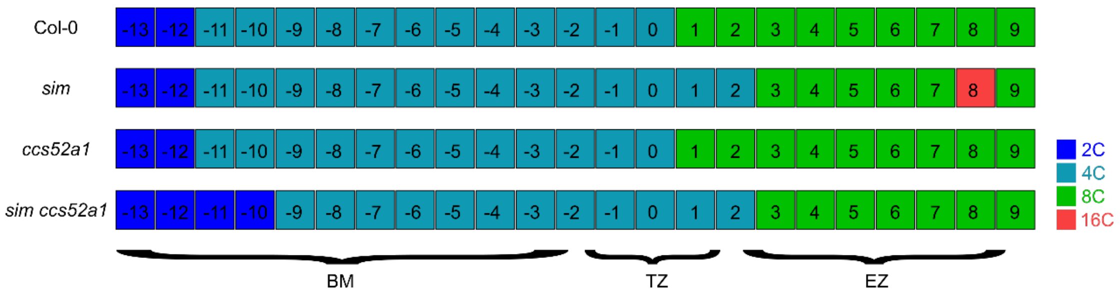


**Supplementary Figure S2.** ***SIM* and *CCS52A1* only marginally control the ploidy level of N epidermal cells.** Mapping of N cell DNA ploidy levels along the root longitudinal axis in WT (Col-0) and *sim*, *ccs52a1*, and *sim ccs52a1* mutants based on DAPI staining. Data sourced from 4 to 5 independent roots of each genotype.

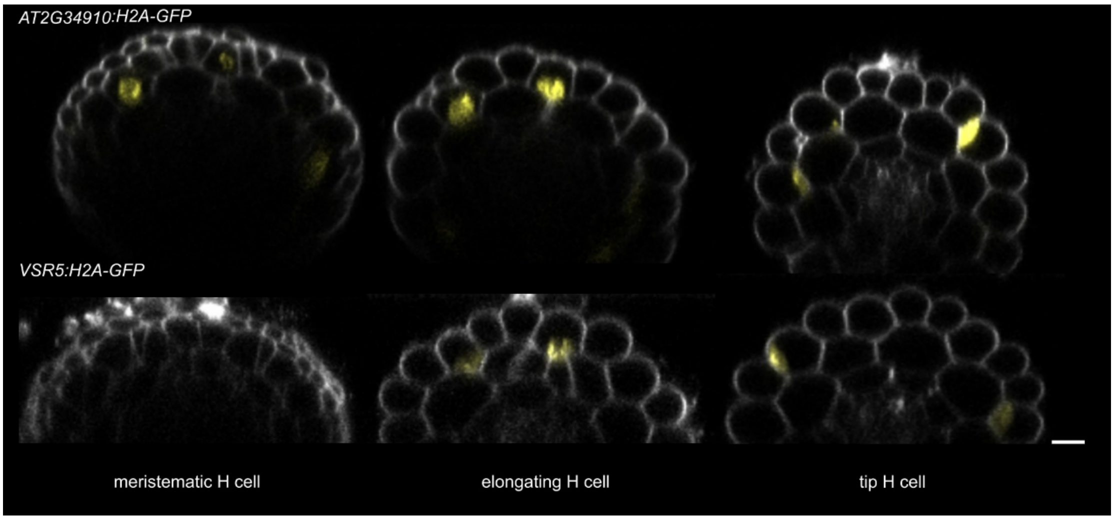


**Supplementary Figure S3. Expression patterns of *AT2G34910:H2A-GFP* (top row) and *VSR5:H2A-GFP* (bottom row) marker lines in the root.** Representative confocal root cross images in the meristematic H (left), elongation H (middle), and tip H cells (right). Images show a GFP (yellow) signal in the H cells. Cell walls were visualized by propidium staining (white). Scale bars, 20 µm.

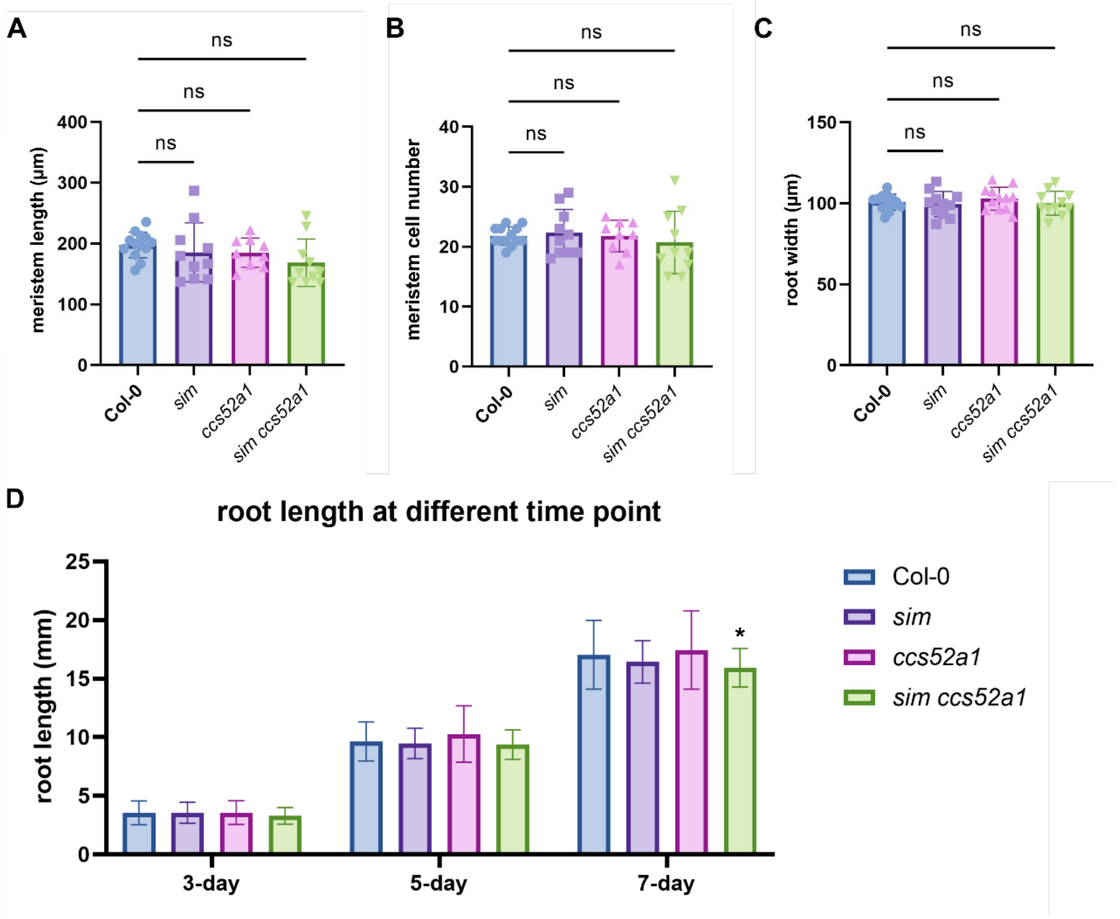


**Supplementary Figure S4. Primary root phenotyping of endocycle mutants in this study. A-C**, Meristem length (A), meristem cell number (B) and root width (C) of WT (Col-0) and *sim*, *ccs52a1*, and *sim ccs52a1* mutants. Analysis was conducted on at least 8 independent roots per genotype. Data are represented as means $\pm$ SD. Significance was analyzed by parametric one-way ANOVA followed by Dunnett's test post hoc test. **D**, Root length of WT and mutants at indicated time points. At least 30 independent roots of each genotype were examined, and the significant differences were analyzed by two-way ANOVA followed by Dunnett's post-hoc test. Significant differences are indicated by an asterisk compared to the control ($p < 0.05$).

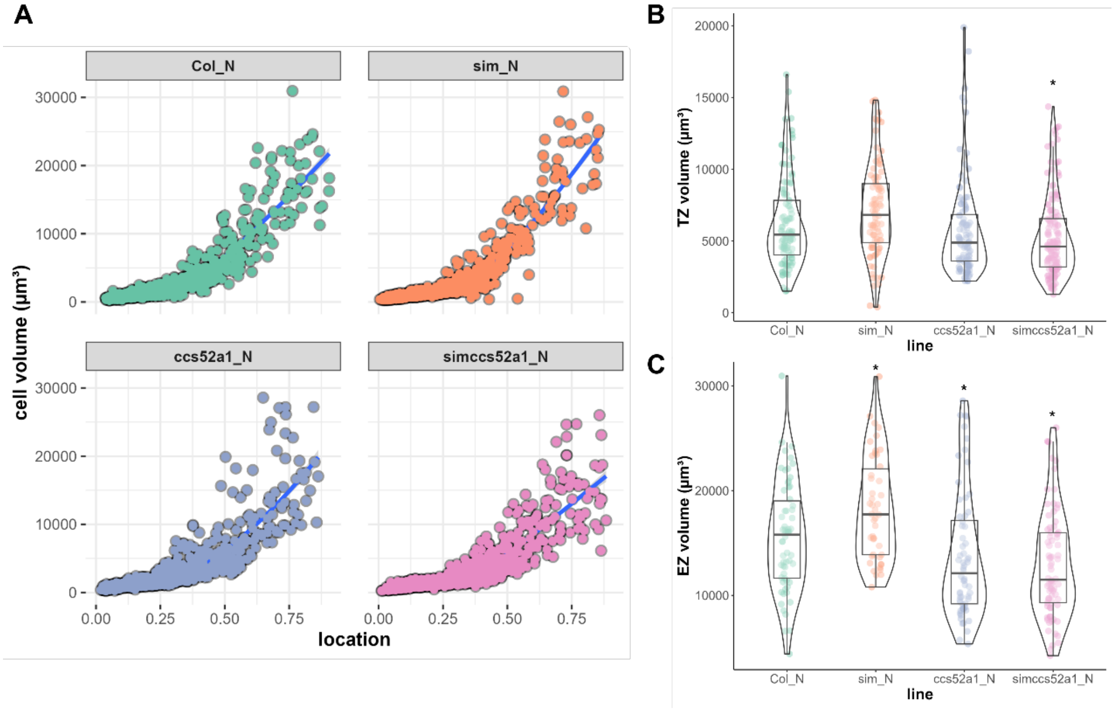


**Supplementary Figure S5. Quantification of the effects of *SIM* and *CCS52A1* on N cell volume in the primary root. A**, Quantification and mapping of the N cell volume along the root longitudinal axis from QC (0.0 point) to the fifth/sixth elongated cell (1.0 point) in the primary roots of WT (Col-0) and *sim*, *ccs52a1*, and *sim ccs52a1* mutants. Over 600 cells among 3 independent roots for each genotype were measured. **B-C**, Cell volumes of WT and mutants N cells in the TZ (B) and EZ (C). Asterisks indicate significant differences (adjusted $p < 0.05$) determined by nonparametric one-way ANOVA followed by Dunnett's post hoc test, compared to WT ($n \geq 40$ per genotype).

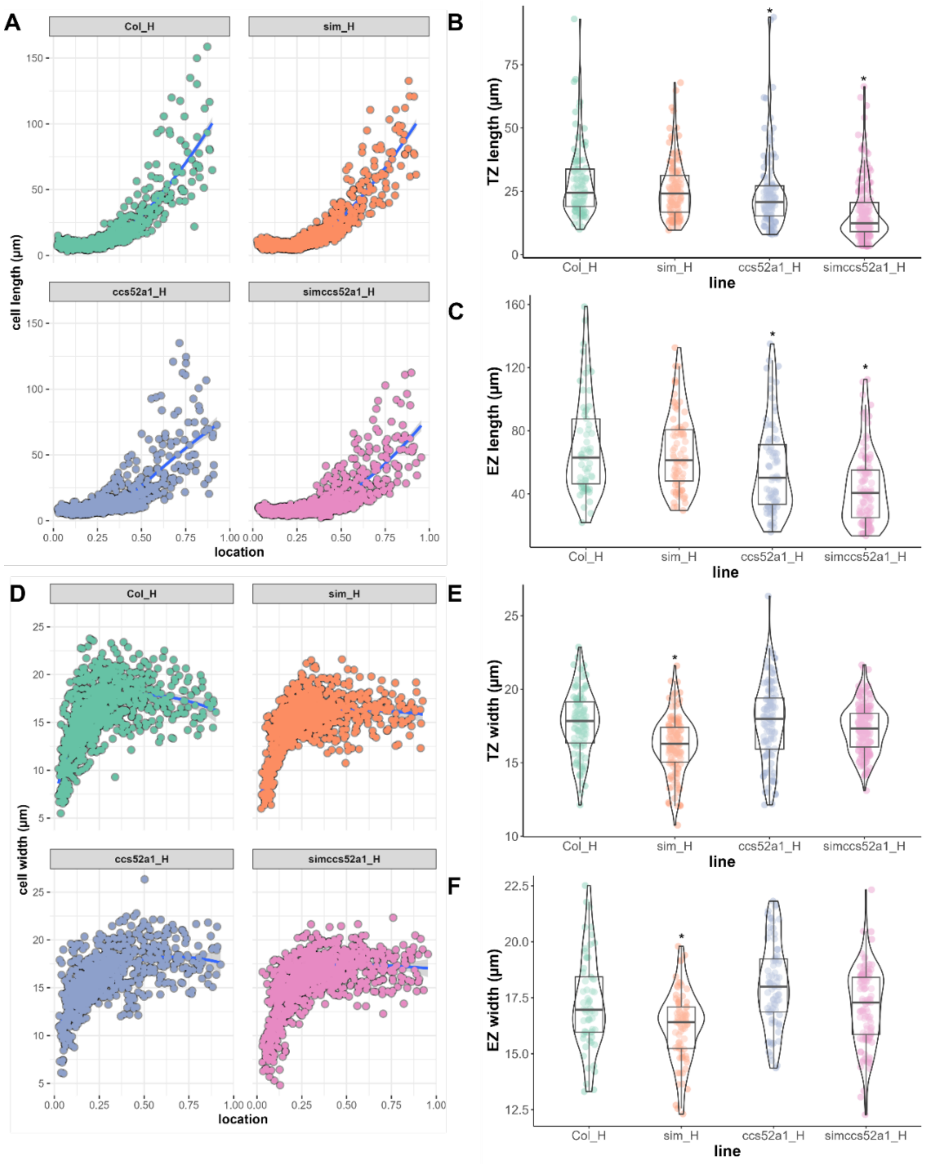


**Supplementary Figure S6. Quantification of effects of *SIM* and *CCS52A1* on the length and width of H cells in the primary root tip. A and D**, Quantification and mapping of the H cell length (A) and width (D) along the root longitudinal axis from QC (0.0 point) to the fifth/sixth elongated cell (1.0 point) in the primary roots of WT (Col-0) and *sim*, *ccs52a1*, and *sim ccs52a1* mutants. **B and C**, H cell length of WT and mutants in the TZ (B) and EZ (C). **E and F**, H cell width of WT and mutants in the TZ (E) and EZ (F). Over 600 cells across 3 independent roots per genotype were measured. Significant differences compared to the control were analyzed by nonparametric one-way ANOVA followed by Dunnett's post hoc test (adjusted $p < 0.05$, indicated by asterisks).

**Table S1. Arabidopsis lines used in this study**

| Line | Description |
|---|---|
| ***sim*** | Bhosale et al., 2018 |
| ***ccs52a1-1*** | Salk_083656 |
| ***sim ccs52a1*** | Crossed line |
| ***SIM:GUS*** | Bhosale et al., 2018 |
| ***CCS52A1:CCS52A1-GUS*** | Willems et al., 2020 |
| ***AT2G34910:H2A-GFP*** | This study |
| ***VSR5:H2A-GFP*** | This study |
| ***AT3G09330:H2A-GFP*** | This study |

**Table S2. Sequences of promoters used in root hair-specific marker lines.**

| Promoter | Length (bp) | Description | Sequence (5' to 3') |
|---|---|---|---|
| **AT2G34910** | 2148 | Forward | AGAAGTGAAGCTTGGTCTCAACCTTAAGGTCAGGGCTGGAAACA |
| | | Reverse | AGGGCGAGAATTCGGTCTCATGTTAACAAAGTCAGAAATCTTAA |
| **AT1G27740** | 1434 | Forward | AGAAGTGAAGCTTGGTCTCAACCTATACATGCATGAATAACGAA |
| | | Reverse | AGGGCGAGAATTCGGTCTCATGTTTAACTTATTTGGATGAAGCT |
| **VSR5; AT2G34940** | 1550 | Forward | AGAAGTGAAGCTTGGTCTCAACCTGGAATCTCGCCTTGTAGTGC |
| | | Reverse | AGGGCGAGAATTCGGTCTCATGTTTTTCGACAAAAGAATCTCT |

The red letters represent the overhang with module A-B entry vectors.
